# Precision as a metric for acoustic survey design using occupancy or spatial capture-recapture

**DOI:** 10.1101/2020.09.30.321521

**Authors:** Julius Juodakis, Isabel Castro, Stephen Marsland

## Abstract

Passive acoustic surveys provide a convenient and cost-effective way to monitor animal populations. Methods for conducting and analysing such surveys, especially for performing automated call recognition from sound recordings, are undergoing rapid development. However, no standard metric exists to evaluate the proposed changes. Furthermore, most metrics that are currently used are specific to a single stage of the survey workflow, and therefore may not reflect the overall effects of a design choice.

Here, we attempt to define and evaluate the effectiveness of surveys conducted in two common frameworks of population inference – occupancy modelling and spatially explicit capture-recapture (SCR). Specifically, we investigate precision (standard error of the final estimate) as a possible metric of survey performance, but we show that it does not lead to generally optimal designs in occupancy modelling. In contrast, precision of the SCR density estimate can be optimised with fewer experiment-specific parameters. We illustrate these issues using simulations.

We further demonstrate how SCR precision can be used to evaluate design choices on a field survey of little spotted kiwi (*Apteryx owenii*). We show that precision correctly measures tradeoffs involving sampling effort. As a case study, we compare automated call recognition software with human annotations. The proposed metric captured the tradeoff between missed calls (8% loss of precision when using the software) and faster data throughput (60% gain), while common metrics based on per-second agreement failed to identify optimal improvements and could be inflated by deleting data.

Due to the flexibility of SCR framework, the approach presented here can be applied to a wide range of different survey designs. As the precision is directly related to the power of detecting temporal trends or other effects in the subsequent inference, this metric evaluates design choices at the application level, and can capture tradeoffs that are missed by stage-specific metrics, thus enabling reliable comparison between different experimental designs and analysis methods.

## 1 Introduction

Sound has been used to monitor vocalising animal populations for many years. Vocal cues are counted in field surveys, and the counts are used directly as an index of population size, or as input to more complex models for inferring species spatial distribution, temporal trends, community diversity, or other properties (see review in Gibb et al. (2018)). Over the past two decades, this use of acoustics has been changed drastically by the development of autonomous recording units, ARUs (Brandes, 2008; Blumstein et al., 2011; Shonfield and Bayne, 2017; Darras et al., 2019).

In their most basic usage, the recorders act as a replacement for field observers in order to collect survey data at a lower cost (Lemckert et al., 2005; Williams et al., 2018), but they have great potential, opening up entirely new monitoring possibilities, such as analysing the ultrasound range for bats (Sugai et al., 2018), localising the calling animals (Collier et al., 2010), or using automated sound detectors to speed up processing (Potamitis et al., 2014). This has sparked various developments in acoustic survey methodology: studies have investigated the optimal number and placement of recorders (e.g., Pérez-Granados et al. (2018); see also the review in Sugai et al. (2019)), compared hardware (Rempel et al., 2013; Pérez-Granados et al., 2019), and optimised recording time and duration (Cook and Hartley, 2018; Hagens et al., 2018; Pérez-Granados et al., 2018). The large amounts of data generated by ARUs has also prompted interest in algorithms to perform call denoising, detection and recognition (Acevedo et al., 2009; Priyadarshani et al., 2018b; Stowell et al., 2018).

However, no unified metric to evaluate these various developments exists. For example, denoising success is usually measured by signal-to-noise ratio (Priyadarshani et al., 2016) or by successful species classification after denoising (Connor et al., 2012). In turn, the species classification methods are evaluated by measuring their agreement with manual annotations, i.e., area under the receiver operating characteristic curve (AUC), accuracy, sensitivity, or specificity (see Methods or Knight et al. (2017) for definitions). These measures can vary hugely between datasets: methods with claimed accuracy of >90% can turn out to be only 40% accurate on a different dataset (Priyadarshani et al., 2018b). Further complications arise because the target is not inherently binary – agreement can be measured per time unit, at the syllable level, call level, or file level, with different results (cf. Acevedo et al. (2009); Swiston and Mennill (2009); Towsey et al. (2012); Potamitis et al. (2014)). In fact, a perfect match with human annotations may not be achievable at all, given that even experts do not always agree (Mortimer and Greene, 2017). This lack of standardisation hampers any attempt to compare or improve the methods.

Another issue that complicates survey method comparison is that improvement in one particular part of the process does not necessary correspond to more accurate or reliable monitoring overall. Compared to field observations, sound recorders have lower detection radius (Yip et al., 2017) and do not capture visual cues, but having recorded data provides many other benefits (Darras et al., 2019). Automated methods for annotation unavoidably miss some calls and likely introduce false positives, in return for faster processing, while additional data sources, such as radio tracking, may come at an extra cost and require sacrifices in sample size. These tradeoffs are not captured by stage-specific metrics (Knight et al., 2017). As summarised in Blumstein et al. (2011), “there is a growing need for a common framework in which to develop, run and fully evaluate new bioacoustic recognition systems. Such a framework would include standard performance metrics”.

For single-species, single-population surveys, which estimate one quantity of ecological interest, a natural choice of metric is the final precision (i.e., the variance) of this estimate. This metric corresponds directly with the power of making subsequent inferences, such as detecting abundance changes, tracking colonisation, or identifying biological factors influencing the species distribution (Stevenson et al., 2015; Metcalf et al., 2019). Thus, it allows the evaluation of changes to any part of the survey protocol at the final stage of ecological application. We focus on two popular statistical survey frameworks: occupancy models (which aim to estimate the fraction of area occupied by the population) and spatial capture-recapture (SCR), which estimates the density of animals in the survey area; these are summarised in the Methods section. We investigate how the variance of the density or occupancy estimate from such models can be used for evaluating changes in survey design.

Specifically, we describe the concept of optimal design in these two frameworks, based on the precision of the final estimate. In each case, we investigate how this optimum depends on unobserved parameters, using simulations and approximations recently developed by Efford and Boulanger (2019). We then focus on SCR and use field data from a bird survey to demonstrate how the precision of density estimate can be used to evaluate tradeoffs involving sample size, and apply this framework to compare automatic call detection software with human listeners. We show that commonly-used metrics of classification accuracy can support detrimental design choices. Finally, we discuss the remaining challenges that need to be overcome to enable standardised survey evaluation.

## 2 Methods

### 2.1 Occupancy modelling

Occupancy models (MacKenzie et al., 2002) seek to determine what fraction Ψ of the survey area is occupied by the species of interest, based on *S* monitoring sites placed in the area. (Each site corresponds to a sensor such as a recorder.) As the detection is not perfect, an estimate of the probability *p* that a species is detected in a site where it is present is needed; this can be interpreted as the sensitivity of the detector. To obtain this estimate, the survey is repeated *K* times. Then, denoting by *s*_*k*_ the number of sites where the species was detected on occasion *k*, and 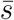 the number of sites where the species was detected on at least one occasion (i.e., was present, assuming no false positives), the occupancy model corresponds to the following likelihood:

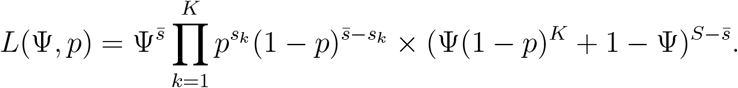

Various extensions to this model have been proposed, allowing, for example, different numbers of surveys at each site, and site-specific detection covariates (MacKenzie et al., 2002; Mackenzie and Royle, 2005; Clement, 2016).

The ultimate aim of this framework is to enable inference about the occupancy parameter Ψ, hence a natural metric for evaluating survey efficacy is the sampling variance of this parameter, 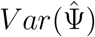 (throughout, we use 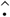 to denote statistics based on samples). The researcher can then choose the experimental design that minimises this variance, and thus maximises the power for subsequent inference, while keeping the estimate unbiased. This approach is well-known in experimental design (Atkinson and Donev, 1992), and in occupancy modelling in particular (Mackenzie and Royle, 2005; Clement, 2016; Gálvez et al., 2016).

Formally, let all parameters that describe the study design comprise a vector ***m*** ∈ *M*. Optimal design for occupancy models can then be defined as:

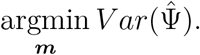

The parameter vector includes *S* and *K* that describe the sampling performed, and also less obvious design attributes that impact the result, such as sensor detection distance and performance of the call recogniser. The space *M* is constrained by the limited experimental effort or technology tradeoffs; occupancy designs are usually analysed assuming fixed total effort *SK* (Mackenzie and Royle, 2005). The detection probability can be separated into a detectability component *p*_*d*_ and an availability component *p*_*a*_, so that *p* = *p*_*d*_*p*_*a*_ (Riddle et al., 2010). Since *p*_*d*_ can be controlled, it is included in ***m***, but *p*_*a*_ is represents the behavioural properties of the species and so is outside the control of the investigator.

As the true values of the ‘nature’ parameters Ψ and *p*_*a*_ are not known in advance, ideally the optimal design should be insensitive to them, i.e.,

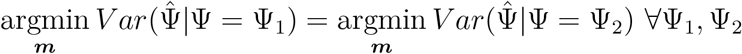

which is true iff:

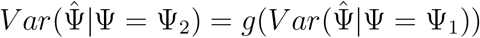

for some strictly monotonic *g*(*x*) *>* 0.

However, the occupancy model does not allow the effects of these parameters to be isolated as monotonic functions. With constant *p*_*a*_, *p*_*d*_, *SK*:

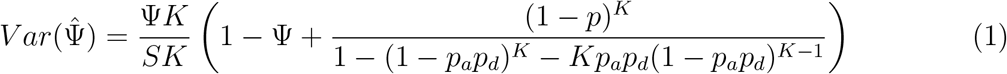

and so:

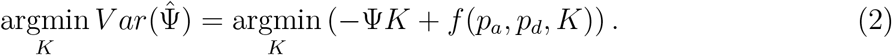

for some appropriate function *f* (*·*).

Both terms in the right hand side expression contain *K* and are non-negligible, unless *p*_*a*_*p*_*d*_ *→* 1, so the optimal choice of *K* depends on the true value of Ψ. Similar results hold for other parameters. For example, if there is a tradeoff between improved detection and more sampling effort (so that *SKp*_*d*_ is constant), then both *K* and *p*_*d*_ will be involved in both terms, meaning that there is no general solution without a priori knowledge of their values.

### 2.2 Spatial capture-recapture

Spatially explicit capture-recapture (SCR, sometimes also abbreviated to SECR) is an alternative framework for detection data analysis (Efford, 2004; Efford et al., 2009; Dawson and Efford, 2009; Borchers et al., 2015), which aims to take into account distance between the animal and the detector. The main inference target is the animal density *D*, which is assumed to be uniform over the survey area and time, or have some parameterised functional form. Animal home range centroids or other location descriptors ****x**** are assumed to be realisations of a Poisson process with intensity equal to *D*. A detection function *p*(*d*|***θ***) describes the probability of detecting animals located at distance *d* from a sensor given parameters ****θ****. The observed data is recorded as a binary capture history over *K* sampling occasions (e.g., days), *S* sensors, and *N* individual detected animals: **Ω**= {*ω*_*skn*_} where each *ω*_*skn*_ = 1 if sensor *s* ∈ *S* detected the particular animal *n* ∈ *N* during occasion *k* ∈ *K*.

The corresponding likelihood is of the form:

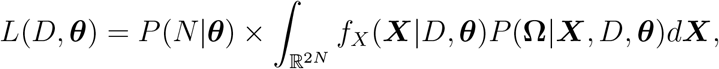

where ****X**** denotes space (so *f*_*X*_ is the pdf of animal locations) and ****θ**** contains the parameters for the chosen detection function. The basic SCR form can be extended to allow non-homogeneous density, customized detection probability function, or additional information collected by the sensors; it allows great flexibility to analyse diverse experimental designs, and incorporate environment and behaviour covariates (Borchers and Efford, 2008; Efford et al., 2009; Dawson and Efford, 2009; Reich and Gardner, 2014; Stevenson et al., 2015; Borchers et al., 2015; Boulanger et al., 2018).

For analysing acoustic surveys, the above model is not immediately applicable. The basic operational unit of acoustic SCR is a call (i.e. {*ω*_*is*_} is 1 if call *i* is detected on recorder *s*). Call locations are considered dependent, and bootstrap procedures are required to estimate sampling variances of the estimates (Stevenson et al., 2015). The parameter *D* then corresponds to call density, and turning this into the underlying animal density *D*_*a*_ requires either the ability to identify individuals in an unsupervised way, or prior knowledge of their call rate *μ*. The first is generally extremely difficult, and the second may well be hard to identify (see e.g., Digby (2013) for kiwi, the species we focus on in the experimental section). However, even if the true call rate *μ* was known, the animal density would be estimated as 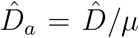 (Stevenson et al., 2015). Therefore, call rate comparisons can be used to test biological hypotheses about population changes, providing that the underlying call rate remains unchanged.

Precision of estimates has been used occasionally for SCR study design (e.g., Sun et al. (2014)), but without investigating the theoretical or empirical grounds for using it – likely because no closed form expression for the variance of the estimates was available. For standard SCR, Efford and Boulanger (2019) recently derived an approximation to the sampling variance of 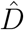, assuming Poisson-distributed number of detected animals *N*, and number of recapture events *R*, as:

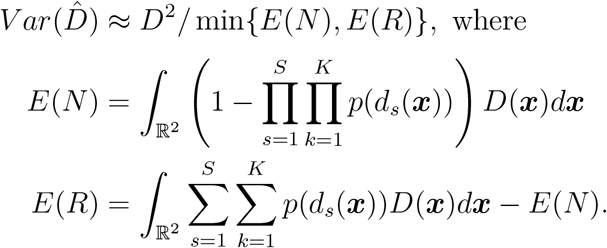

Here *S* and *K* are the number of sensors and survey occasions, respectively, as previously, while *d*_*s*_(****x****) denotes the distance between sensor *s* and animal ****x****, and *p*(*·*) is the detection function.

Acoustic surveys are usually analysed as single-occasion data (*K* = 1), conducted for *τ* time units, and the detections (*N* and *R*) are calls emitted by each animal at a fixed rate *μ* (Stevenson et al., 2015). To apply the above equations to such analysis, we assume that the density (of calls) is homogeneous and equal to *μD*_*a*_, and obtain:

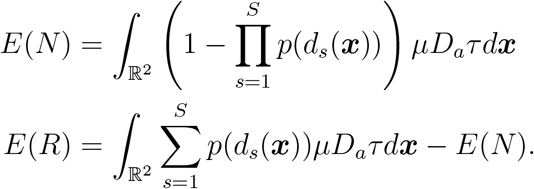

The optimisation task can then be written as:

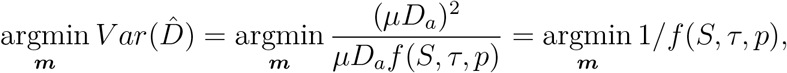

and so the optimal design ****m**** = (*S, τ, p*) is invariant to the true density *D* or call rate *μ* (or any other parameters that have a constant and uniform effect on the cue density). Thus, in this respect, the design for acoustic surveys in the SCR framework can be optimised without prior knowledge of behavioural parameters.

### 2.3 Simulations

We used numeric examples to illustrate the different effects of parameters within occupancy studies and SCR models.

To illustrate that the optimal design for occupancy studies depends on the values of population-specific parameters, we calculated the standard error of 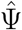 over several values of *p*_*a*_, *p*_*d*_, Ψ and *K*. (Recall that *p*_*d*_ is meant to capture the effect of design choices, such as detection thresholding, while *p*_*a*_ cannot be manipulated by the observer.) We assumed fixed sampling effort of 24 sites, i.e. *S* = 24/*K*, and then simply calculated the variance using Equation (1).

As our results concerning SCR rely on the approximate variance formula, as well as independent call locations, we verified that they are valid for acoustic surveys using simulation. We used the ascr package to generate binary detection histories, as outlined in Stevenson et al. (2015). In summary, the user defines a survey by providing detector positions and an area for integration (we used a 3 × 3, 4 × 4 or 5 × 5 grid of detectors spaced at 200 m, surrounded by a 200 m buffer). Call locations are then simulated in this area. To generate ‘observed’ detection histories at each detector, `ascr` filters these call locations based on detection parameters and distance to each detector. We used a half-normal detection function 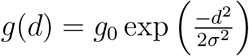. In practice, different hardware, call recognition algorithms or detection thresholds would have an effect on the detection probability, so the study design consisted of detection parameters *g*_0_*, σ*, and number of detectors *S*. We did not vary the survey duration *τ* in the simulation, since as the capture histories do not include actual call times, a longer survey at smaller call rate would be indistinguishable from a shorter survey with higher call rate.

We initially modelled the call locations as independent, i.e., as realisations of a spatial Poisson process with intensity equal to the call density *D* and repeated simulations for several values of *D* between 100 and 500. For each tested combination of design parameters and *D* values, 100 survey datasets were generated, an SCR model fitted to each dataset, and the observed standard deviation of 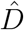 used to estimate the corresponding standard error. Relative bias was estimated as.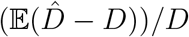.

Since many calls can come from the same bird, or from a group of birds calling in the same place (such as alarm calls, territorial warnings, or duets), the call locations may be dependent in practice. We therefore conducted a similar set of simulations to test the consequences of violating this assumption. In `ascr`, this is implemented by simulating the locations of animals from a process with intensity equal to *D*_*a*_, and repeating calls at each location *μ* times. Several values of *μ* were tested with a fixed call density *D* = 300, 100 replicates simulated, and each detection history analysed in a similar manner to the independent case.

### 2.4 Kiwi survey data

To show how SCR can be used to evaluate design tradeoffs in a practical setting, we conducted an acoustic survey of kiwi pukupuku (little spotted kiwi, *Apteryx owenii*) in Zealandia, a 225 ha wildlife sanctuary in Wellington, New Zealand. The likely population size is around 200 kiwi, with 40 kiwi translocated to this site in 2001, and approximately 100 estimated in 2010 (Digby, 2013). Kiwi pukupuku has a life expectancy of >30 years; once individuals become adult, mortality is low, and pairs produce a 1-2 egg clutch each year (Colbourne, 1992; Robertson and Colbourne, 2004; Jolly, 1989).

This species is nocturnal, and both sexes call intermittently throughout the night, often in duets; male and female calls are different. The calls consist of many repeated syllables slowly rising in pitch, with fundamental frequency around 2-3 kHz for males and 1-2 kHz for females (Supplementary Figure S2). The number of syllables and exact pitch vary between calls. Further details on kiwi acoustics can be found in Digby (2013) and Digby et al. (2014). There are other night callers at the study site, some of which can be confused with the kiwi pukupuku, particularly some calls of the ruru (*Ninox novaeseelandiae*, a native owl).

We used data collected by 7 autonomous sound recorders (Song Meter SM2, Wildlife Acoustics Inc., USA) over two nights in 2018, Oct 6 18:00–Oct 7 06:00 and Oct 7 18:00–Oct 8 06:00. The recorders were attached to trees at about 1.5 m height and spread to cover an area of 600 × 800 m (Supplementary Figure S1), with an average spacing of about 300 m, a distance chosen based on rough estimates of call propagation distance. Recorder positions were projected to metres easting and northing using Universal Transverse Mercator (UTM) zone 60G in R package `rgdal`. In total, 10 080 minutes of audio was recorded across seven recorders.

Data was processed in the AviaNZ software (Marsland et al., 2019). In the automated process, the software was first trained on a small amount of manually annotated data from the same location, collected in September 2018. During this process (see Marsland et al. (2019) for detailed description), the sound is decomposed into a wavelet representation: this basis of compact functions allows to quantify the energy in different frequency bands (nodes) at discrete time points. The software identifies an optimal set of wavelet nodes for automated call detection. We trained two filters for the kiwi, using ‘high’ and ‘low’ (fundamental frequency above or below 2 kHz, approximately separating males and females) calls separately. Detection thresholds were set to obtain 90% sensitivity on the training data.

The two nights studied (Oct 6 and Oct 7) were automatically processed using these filters: as in training, the software reconstructed the signal from selected wavelet nodes and marked putative calls when high energy in the reconstructed signal was detected. This approach is optimised to avoid false negatives, which can result in a high number of false positive marks from the software; this is a user-controllable parameter, and could be chosen to avoid false positives, albeit at the cost of missing many actual calls, and so we chose the former for the experiments reported here. All the potential ‘calls’ detected were reviewed by a human observer, to ensure that final detections contained only true positives.

The audio data, automatic recognizer settings used for detection in this paper, and AviaNZ format annotations are available at https://doi.org/10.5061/dryad.m70p89d (Oct 6th data) and http://doi.org/10.5281/zenodo.4057094 (Oct 7th data). The AviaNZ software is available at http://www.avianz.net.

### 2.5 Survey data analysis

Acoustic SCR models were applied to the kiwi survey data in several experimental settings. In all analyses, we used standard binary capture histories across the 7 recorders, labelled as shown in Supplementary Figure S1. In such cases, the SCR detection function is estimated only from the frequency of combined detections across multiple recorders, which requires reliable identification of when two or more annotations on different recorders represent the same call. To adjust for recorder clock drift, we measured how many 1-second blocks had calls present on two neighbouring recorders, and adjusted the clocks in 1-second steps until this overlap was maximised (Supplementary Figure S3); we visually inspected the spectrograms to confirm that most overlaps after adjustment reflected the same call (Supplementary Figure S4). For some pairs of recorders, no clear adjustment was identified, and most overlapping annotations appeared to be different calls. In effect, the 7 recorders were split into 3 non-overlapping groups, ZB-ZI-ZG; ZE; ZA-ZH-ZJ. Annotations were then converted into a call capture history for each recorder. Annotations separated by <5 seconds were merged into a single call, to avoid double-counting fragmented calls or female responses to male calls.

All statistical analyses were conducted using `R`. For SCR modelling, we used the `ascr` package, which is available from https://github.com/b-steve/ascr. All other code used in the analysis is available at https://github.com/jjuod/acoustic-survey-evaluation.

### 2.6 Measuring survey precision on data subsets

In theory, SCR precision is related to the duration of acoustic survey as the standard error 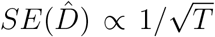. To verify that this holds in practice, we fitted acoustic SCR models to subsamples of the acoustic survey conducted over two nights. We fitted SCR models using automatically annotated (human-reviewed) data from the two nights, or subsets of 75%, 50%, 25%, or 12.5% of it, by using only annotations within the respective fraction of each hour (e.g., for the 25% fraction all annotations outside 18:00-18:15, 19:00-19:15, etc., were ignored).

We used the half-normal function to model detection probability. The area of integration was defined by setting 700 m radius circles around each recorder, as the estimated detection probability in our setup decayed to <0.005 at this distance. The main outputs of interest were the call rate density 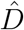 and effective survey area (ESA) in hectares, derived from the detection parameters; the effective detection radius was then calculated as 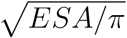. The standard errors were calculated using the Hessian or by bootstrapping – drawing 200 replicate surveys of the same length (Stevenson et al., 2015) – and converted to units of calls/ha/night.

### 2.7 Comparison of human and software call detection

In the next experiment, we used the SCR framework to compare automatic and manual call annotation processes. Manual annotations were made for data from the Oct 6 night, using audio playback and visual inspection of the spectrogram in AviaNZ (Marsland et al., 2019). The processing was split between two observers, with each annotating ca. half of the dataset. Automatic (human-reviewed) annotations for this night were already obtained as described previously. We fitted two SCR models using either the manual or automatic (reviewed) annotations, and calculated the precision of the final density estimate using bootstrap sampling. For comparison, we also calculated metrics based on agreement between automatic and manual annotations. Using 1 s resolution, we calculated the numbers of seconds corresponding to true/false negatives/positives (treating manual annotations as the ground truth), and derived sensitivity (or recall, *TP* /(*TP* + *FN*)), specificity (*TN*/(*TN* + *FP*)), and accuracy ((*TP* + *TN*)/*total*), using standard definitions (Knight et al., 2017; Priyadarshani et al., 2018b). Similar calculations were performed using 15 s windows. In each case, a window that includes one or more annotations is considered a positive for the source of annotations. Finally, we investigated the effect of recorder removal in a similar setup, using data from the night of Oct 6. We excluded two recorders with the highest levels of noise (ZB and ZH) or worst agreement with human annotations (ZA and ZI). SCR models were then re-fitted using the remaining data, and all metrics recalculated.

## 3 Results

### 3.1 The effect of population parameters on experimental design

It is generally known that optimal design for occupancy studies depends on the true occupancy (Mackenzie and Royle, 2005). We replicated this result, with separate availability and detectability components *p*_*a*_ and *p*_*d*_, as shown in Figure 1. For example, designs with higher replication effort (*K* = 6) were optimal when *p*_*a*_ = 0.5, Ψ = 0.9, but they were the worst out of the six tested designs when *p*_*a*_ = 0.9, Ψ = 0.7. Design ranking was not consistent across the other conditions either. A relatively small difference in parameter values was sufficient to cause drastic changes in design ranking.

**Figure 1:**
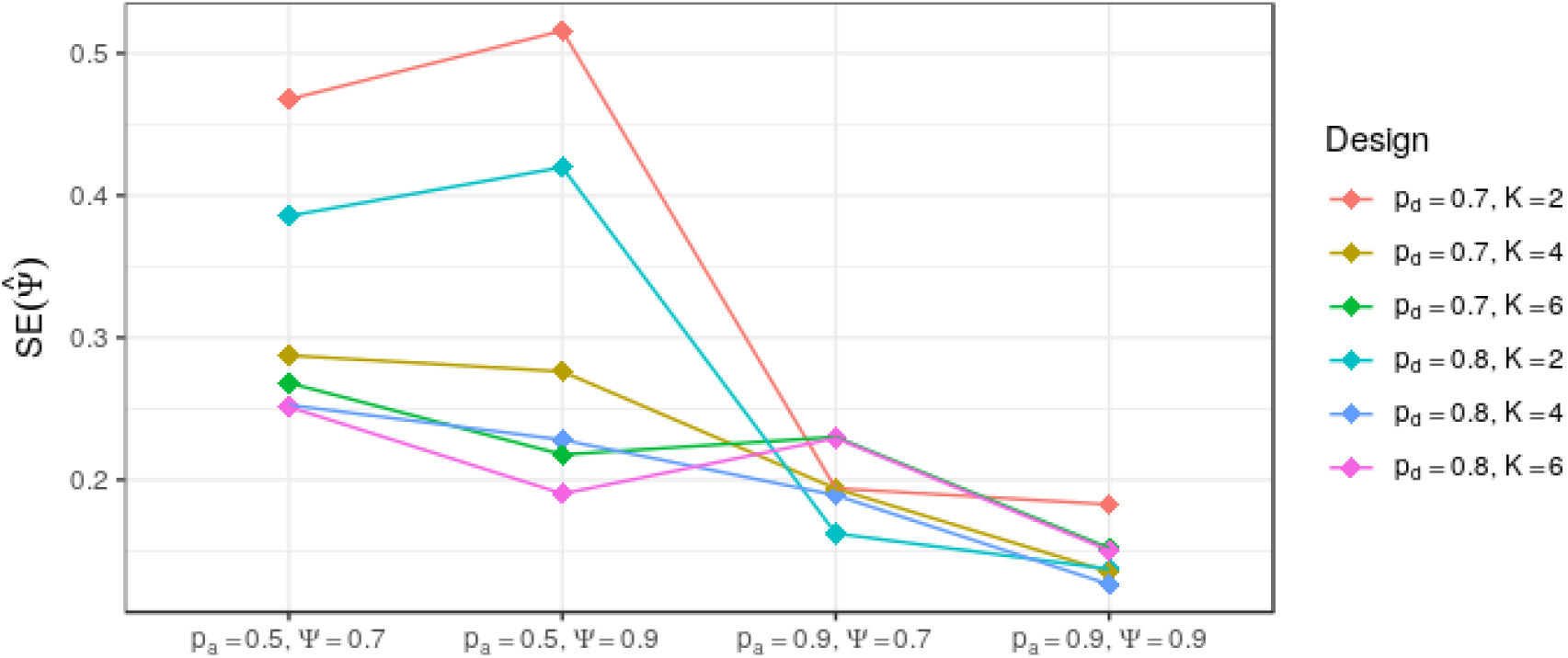
Standard error of the occupancy estimate 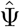 for different combinations of “nature” parameters (availability *p*_*a*_, true occupancy Ψ) and design choices (detectability *p*_*d*_, survey occasions *K*). Units of occupancy are fraction of area occupied (range 0 to 1, true values 0.7 or 0.9). Standard errors calculated analytically, so no certainty bars provided.

In contrast, designs identified as optimal in the SCR framework are expected to remain optimal under a broad range of density values. Figure 2 shows the estimated precision (standard error) obtained under 9 different study designs (represented by different *g*_0_*, σ, S* values). The design with *g*_0_ = 0.8*, σ* = 60, and a 4 × 4 detector grid was optimal under all tested values of *D*; in general, the ranking of designs remained stable across the entire range of *D* values tested. The design with *g*_0_ = 1*, σ* = 30 and a 5 × 5 detector grid was the worst in all conditions, and produced considerably biased estimates (up to 15 % lower; Supplementary Figure S5). All other designs estimated densities within 3% of the true value.

**Figure 2:**
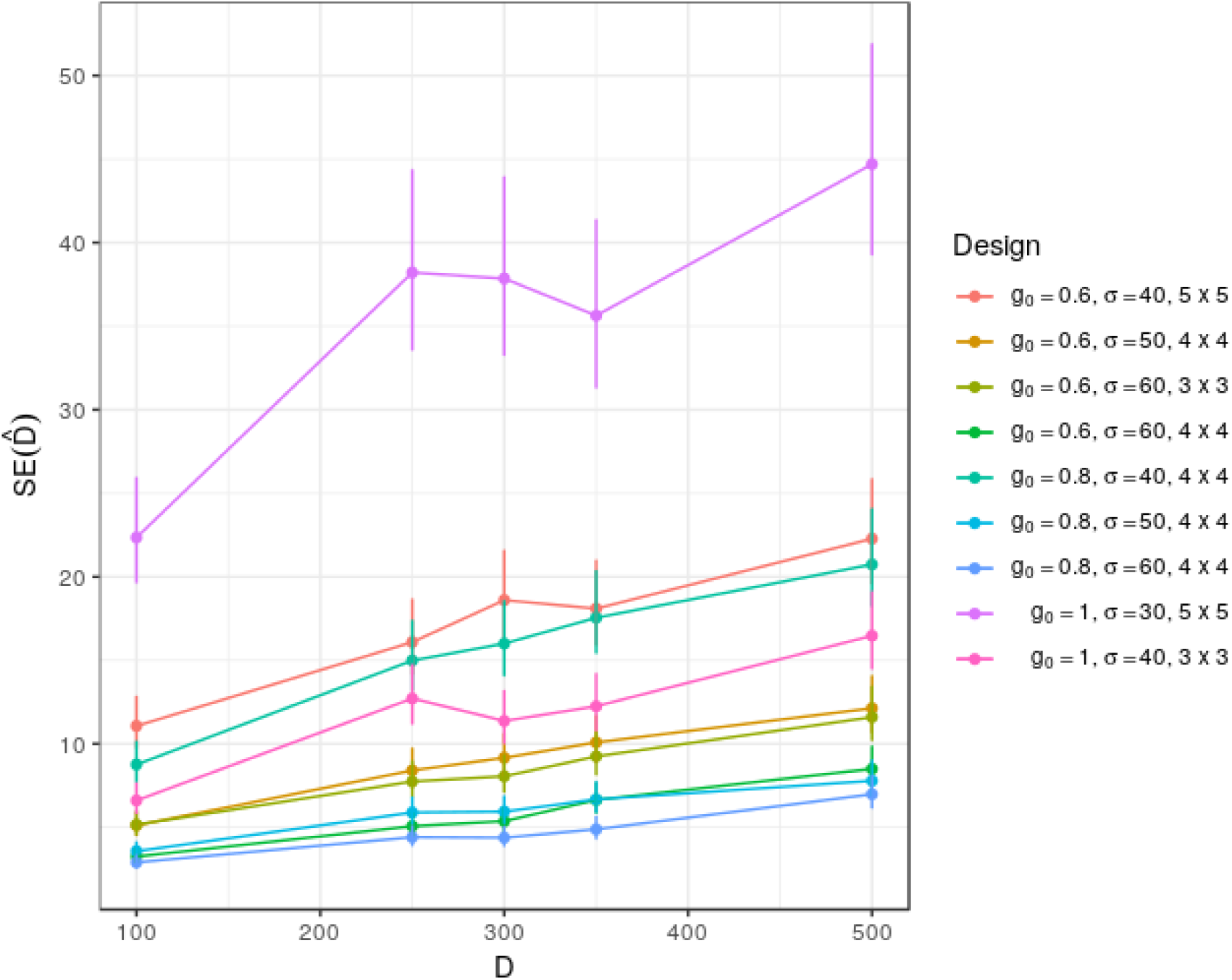
Standard error of SCR density estimate 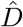 for different values of true density *D* and various design choices. Design choices are represented by different numbers of sensors (*S* in Methods) and different parameters of the detection function (*g*_0_, detection probability at 0 m distance, and *σ*, controling detection decay). For each setting, capture histories were simulated, assuming independent call locations, and density estimated with SCR. Dots are the standard errors based on 100 replicates, and vertical bars indicate 95% confidence intervals based on *χ* ^2^ distribution of the SE. Density units are calls per ha per survey.

Similar results were observed when dependency in call locations was introduced: relative efficiency of study designs for SCR remained stable across various values of call rate *μ* (Figure 3). Some minor deviations were observed, likely attributable to sampling variation in the sampling error itself. Bias was also similar to the independent case, at most 3.4%, apart from the pathological design, which had 5-10% bias (Supplementary Figure S5).

**Figure 3:**
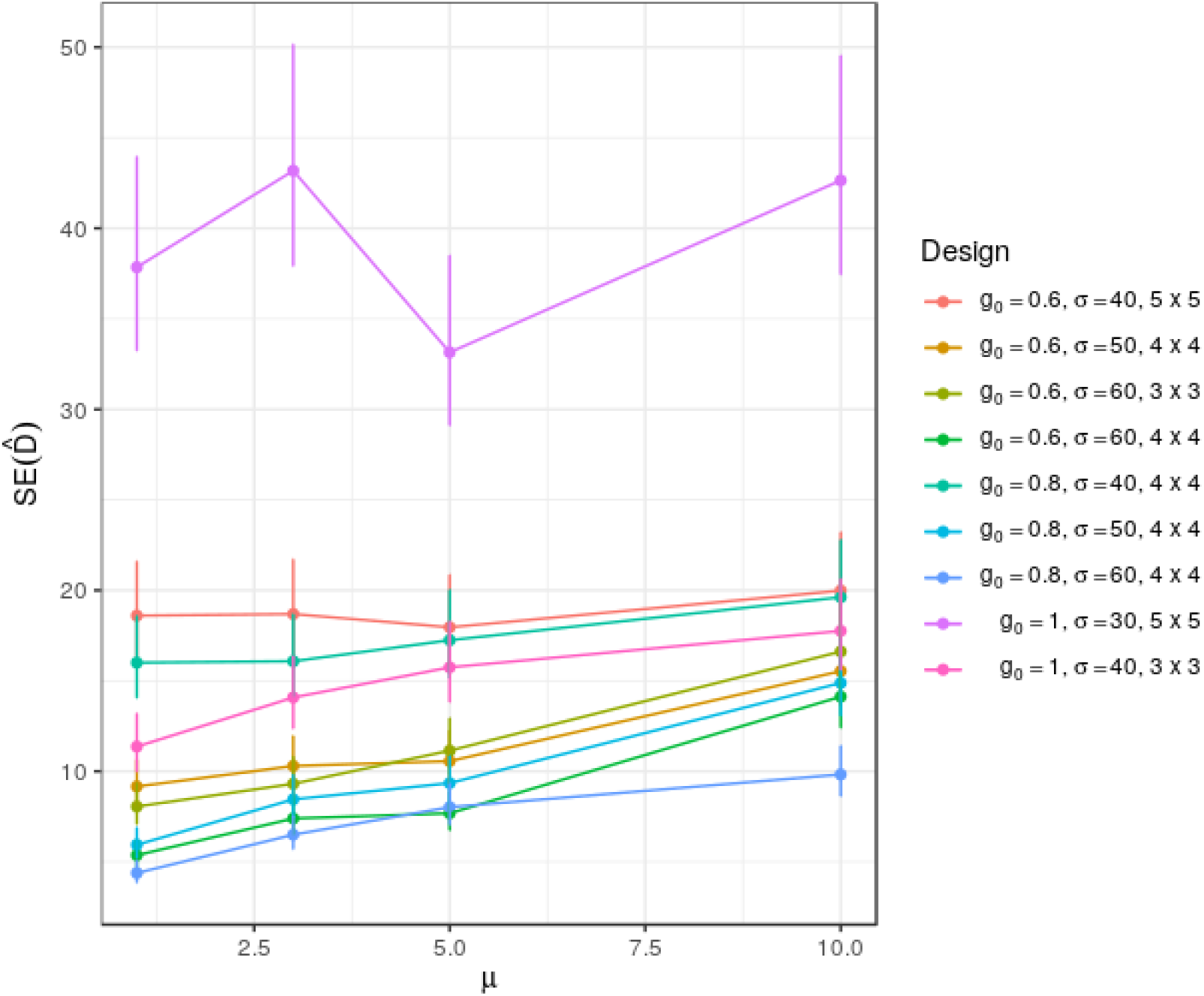
Standard error of SCR density estimate 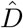 for different values of true call rate *μ* and various design choices. Design choices are represented by different numbers of sensors (*S* in Methods) and different parameters of the detection function (*g*_0_, detection probability at 0 m distance, and *σ*, controling detection decay). Capture histories were simulated for each setting, with total call density equal to 300 and calls repeated *μ* times in each position. Dots are the standard errors based on 100 replicates, and vertical bars indicate 95% confidence intervals based on *χ*^2^ distribution of the SE. Density units are calls per ha per survey.

### 3.2 Precision accounts for changes in sampling effort

We now focus on SCR. In the first experiment based on real data, we investigated how SCR estimates respond to changes in the amount of data processed.

Automatic detection using the AviaNZ software identified 5117 minutes of putative calls, of these 3372 were suggested by the ‘high’ (male) and 1745 by the ‘low’ (female) call filter. Subsequent human review rejected most of these as false positives, leaving 514 min of calls as the final result from this method of processing (370 from ‘high’ and 144 from ‘low’ filter detections).

By fitting an SCR model to this dataset, a call rate density of 2.86 calls per hectare per night was obtained (bootstrapped SE 0.146 calls per hectare per night), based on an effective survey area of 103.4 ha. The detection function indicated that the probability of detecting a call emitted at 200 m distance was 54% (for each recorder), or equivalently the effective detection radius was 256 m.

When the data was subsampled to contain only a fraction of each hour, the observed precision closely matched the predicted square-root relation (Figure 4). The asymptotic and bootstrapped standard errors were very similar, as we did not assume any dependency structure in the call locations. Hence, the precision measure can be immediately translated into sampling effort and power: a researcher wishing to detect two times smaller effects at the same power will need four times more audio data, for example.

**Figure 4:**
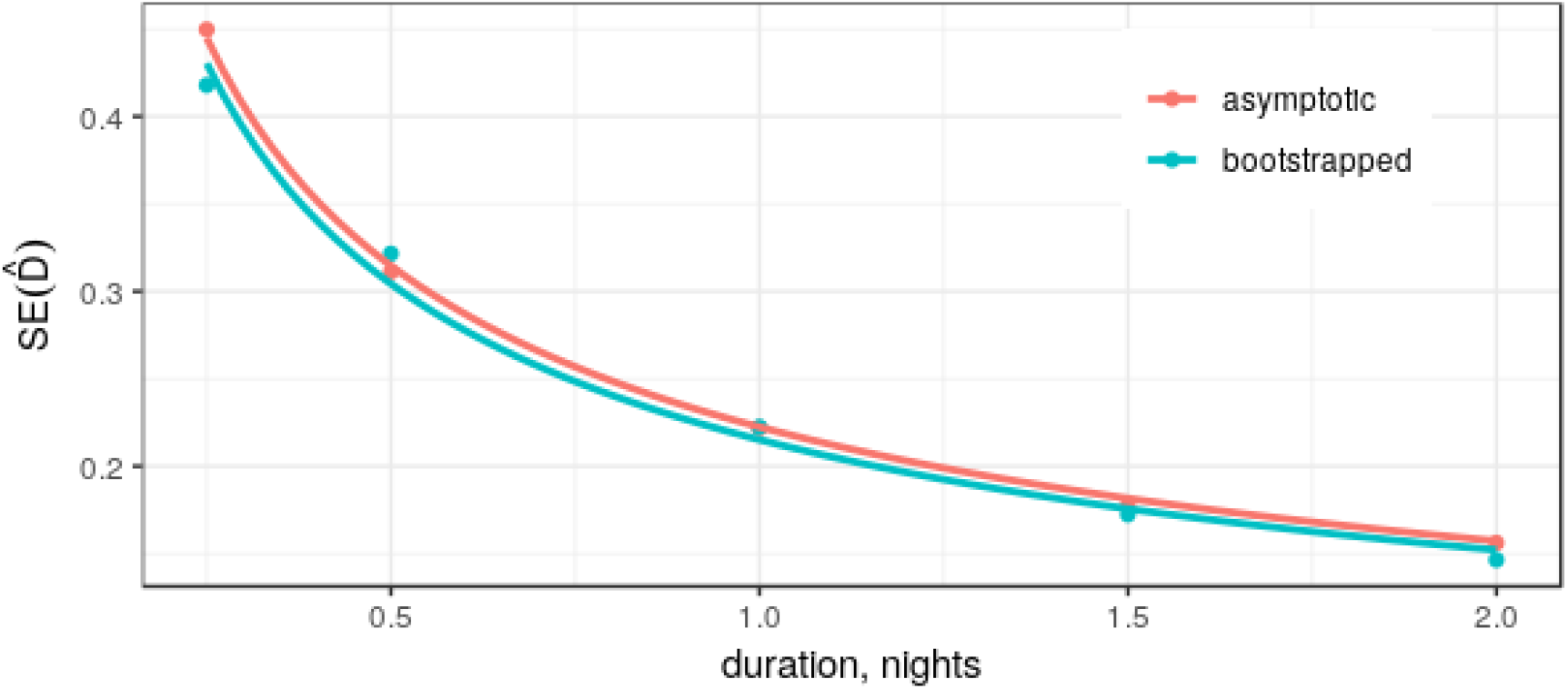
Standard error of estimated call rate density using a full two-night dataset from a kiwi survey (rightmost points), or subsamples of it. Points are measured SEs (obtained from asymptotic approximations or by bootstrapping), the lines are best-fit inverse square-root curves.

### 3.3 Comparing recognition software with human detection

As a case study demonstrating the usage of the proposed metric, we compare the use of automatic call detection software with fully manual processing of recordings. In manual analysis of the first (Oct 6) night of recordings, 915 annotations were made, comprising a total call duration of 232 min (Supplementary Figure S6). Excluding annotations <5 seconds, which usually indicate fragments of a single call, 720 calls remained. Processing 42 hours of audio took about 320 minutes of active user time. The distribution of calls detected by different recorders varied, in particular fewer calls were detected by recorders placed near to a stream (ZE) and on a windy hilltop (ZB).

The automatic processing results (after review) largely matched the manual annotations. Using the human annotation as ground truth, at 1 second resolution, 3.5% of the dataset was marked as positives by both methods, and 94.2% as negatives by both. 1.6% of the recordings were ‘false negatives’, i.e., manually annotated as calls, but missed by the automated detection, and the remaining 0.7% were detected by the automated, but not the manual, processing. Some of these mismatches were due to imprecise annotation boundaries (e.g., automatic detection can miss part of a call, or include a few seconds outside the call, both of which are penalised), others represent genuine missed calls.

Using SCR, we obtained similar density estimates for the two processing methods, with slightly lower precision when calls were marked automatically (Table 1, top rows). Density values were well within one SE of each other, and no systematic difference was apparent (2.93 vs. 2.89 calls per hectare per night). The automatic processing was worse at detecting faint calls, which in SCR corresponds to a smaller survey area (114.3 vs. 99.3 ha) and larger standard error of the estimated density (0.215 vs. 0.227 calls/ha/night). In terms of the detection function, human processing resulted in 61% detection probability at 200 m, in contrast to 54% measured above.

**Table 1:**
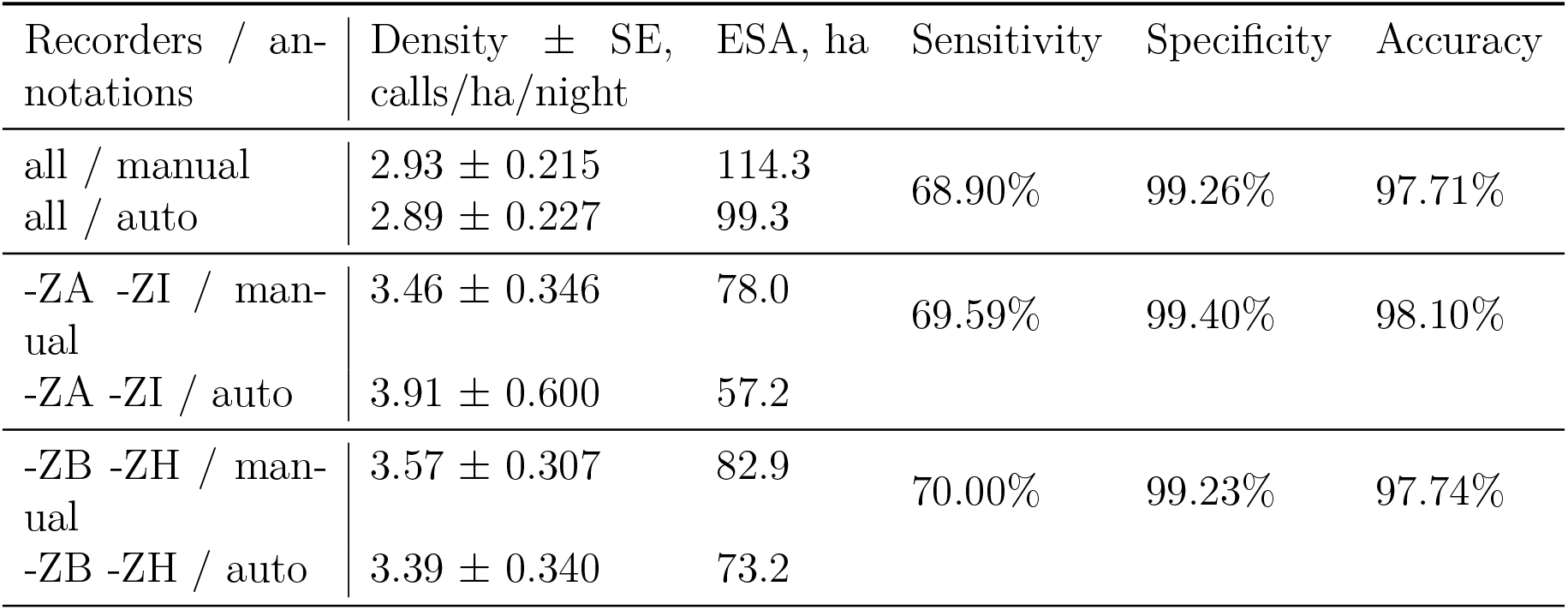
Changes in concordance and SCR measures using all 7 recorders, excluding ZA and ZI, or excluding ZB and ZH. Estimated call rate density, effective survey area (ESA), and their standard errors calculated for manual or automatic annotations separately, using Oct 6 night data. Sensitivity, specificity, accuracy obtained by comparing manual and automatic annotations over 1 second blocks.

However, manual processing was considerably more labour-intensive. Review of software outputs took 132 minutes, making the automated processing with review 2.4 times faster (in user time) than the fully manual workflow. Equivalently, given fixed total effort, 2.4 times more data could be processed in the automated workflow, with almost all user time spent on the review of the detections. Based on Figure 4, this would correspond to a 60% gain in precision, far outweighing the 8% loss due to missed calls.

### 3.4 Standard agreement measures do not reflect optimal design choices

To provide direct evidence of the shortcomings of standard concordance measures, we showed that the match between manual and automatic annotations can be improved by reducing the dataset. When using data from all 7 recorders, with the human annotation viewed as ground truth, the overall sensitivity of the automatic detection method was 68.9%, specificity 99.3%, and accuracy 97.7% (see Table 1, Supplementary Table S1). Recorder ZA had the lowest sensitivity (59.7%), and ZI the lowest accuracy (96.2%); on these grounds, data from these two recorders was excluded, and all measures recalculated using data from the 5 remaining recorders. Despite the obvious loss of information, all agreement measures increased: sensitivity to 69.6%, specificity to 99.4%, accuracy to 98.1%. Similarly, the ZB and ZH recordings were found to have a great deal of wind noise in them. Excluding these instead gave an even larger increase in sensitivity and almost no change in specificity or accuracy. In contrast, data loss decreased the SCR estimate precision in each case: standard errors were increased by 40-75%, while the density estimate itself was increased by about 20%.

We also note that the agreement measures depended on the resolution at which presence/absence was determined. Using larger windows (15 s instead of 1 s), we obtained considerably higher sensitivity for every recorder combination tested (increase to around 75% from 69%), with specificity still remaining above 99% (Supplementary Table S2). A drop in accuracy to 95.4% was observed, as would be expected when the fraction of positives in the ground truth (or true species prevalence) increases.

## 4 Discussion

Many studies over the last two decades have suggested improvements to the methodology of performing and analysing acoustic surveys for conservation purposes. In this paper we have highlighted some of the issues that arise when trying to evaluate such improvements: (1) optimal design in certain models strongly depends on unknown parameters such as the true occupancy, and (2) the success of a single stage of the survey process does not necessarily reflect the overall performance; these can lead to large uncertainty in abundance estimates, with obvious negative consequences.

The former issue is known in the occupancy modelling literature (Mackenzie and Royle, 2005; Gálvez et al., 2016), but the latter has received less attention: for example, call recognition is mainly evaluated using agreement metrics (e.g., Potamitis et al. (2014); Stowell et al. (2018); Metcalf et al. (2019)), and even though some problems stemming from them have been discussed (Knight et al., 2017; Priyadarshani et al., 2018b; Sofaer et al., 2019), no alternatives have been proposed. In the main comparison presented here – manual versus automatic call detection – we have showed that the standard metrics rate the detection software poorly, since they do not account for the gain in processing speed, nor accurately reflect the impact that missed detections have on the final density estimation. This example highlights that detrimental changes to the survey (deleting large portion of data in this case), can produce no effect, or even appear as improvements, on a single stage of the processing pipeline. Similarly, the standard performance measures depended on the measurement resolution, as seen in literature e.g. Metcalf et al. (2019).

We propose that the issues discussed could be alleviated by using the precision of the SCR density estimate as a metric for evaluating acoustic survey methodology. As it has a close relationship with the inferential power, this metric corresponds to the desired ecological outputs at the application level. Our results demonstrate that this metric properly accounted for changes in effective sample size, and correctly identified that automatic processing provides an overall improvement to survey efficiency.

In practice, this framework can be used by conservation managers to compare candidate survey designs using simulated capture histories, or test different processing softwares using real recordings. In our case, the sampling effort is represented by the human listening time, as that is a major bottleneck in current acoustic surveys (Shonfield and Bayne, 2017), but the same approach can be used to find the best experimental design given total budget or other constraints. Cost-benefit analyses of acoustic surveys have been performed previously, but without a general metric they had to assume that all options have equal detection properties (Lemckert et al., 2005; Williams et al., 2018), or otherwise constrain the compared designs, such as setting a fixed study duration and area (Pérez-Granados et al., 2019). The approach presented here allows comparing diverse study designs directly by summarising multiple variables into the final estimate precision, either with occupancy or SCR models. In both cases, the necessary standard error calculations are readily available in the modelling software, but in occupancy studies the optimal choices of experimental design depend on unobserved parameters, and so design tradeoffs need to be identified on a case-by-case basis and thus are not directly applicable to issues such as recogniser algorithm development.

Even in SCR, the optimal design requires some knowledge of the target species ecology. In particular, the need for overlapping detection ranges means that spacing between detectors must be chosen based on the expected call propagation distance (taking into account that imperfect call recognition will reduce this distance). If the recorder grid is concentrated entirely within a single call range, or spaced too widely, resulting in very few ‘recaptured’ calls, the estimated densities will be considerably biased, and this bias is not captured by our metric. However, there is some evidence (Sun et al., 2014) that SCR models are robust to moderate deviations from optimal spacing.

Knowledge of the call rate *μ* is needed to convert the density of calls into actual population counts, and this is rarely straightforward. The call rate of little spotted kiwi, like many other animal and bird species, is affected by factors such as season, temperature, humidity, population density, and even light levels (Keast, 1994; Digby, 2013; Watson et al., 2016). In this study we therefore limited our field data analysis to the density of calls. However, current monitoring protocols (e.g., Colbourne and Digby (2016)) already assume that the call rate is fixed and equal in all surveys compared, to allow inferring population trends from changes in cue count; under the same assumptions, conclusions derived from relative precision of call density still apply to animal density.

An alternative, which merits further research, is the recognition of individuals from their calls, which can be accommodated by the SCR framework (Stevenson et al., 2015). While individual-specific features in kiwi syllables were shown to exist in Digby et al. (2014) and Dent and Molles (2016), determining individual counts in an unsupervised way for large populations will be a much more complicated task (Gibb et al., 2018), although there has been progress in other bird species, see e.g., Puglisi et al. (2004); Ptacek et al. (2015); Stowell et al. (2017).

The main dataset used in this paper comprises recordings over two nights from seven recorders. These recordings were analysed both automatically (with the outputs verified by humans) and manually, with the aim of demonstrating the use of the precision of the SCR density estimate for this change in analysis method. As the two processing choices differed strongly, we found the two nights sufficient to establish the difference, but larger samples would be needed if surveying cryptic species, or in open areas where wind conditions can cause large variability in recording quality (Priyadarshani et al., 2018a).

While our recording period was short, and longer observation across different seasons would be needed to establish the call rate, the results are in line with more complete studies. Approximate kiwi call rates per individual can be obtained from the data in Colbourne and Digby (2016) to be between 0.64–2.8 calls/night for five sites with brown kiwi (*Apteryx mantelli*), and 1.4 for a survey of great spotted kiwi (*Apteryx haastii*). Assuming kiwi density in our study area was about 1 individual/ha (Digby, 2013), a reasonably similar call rate of 2.9 calls/night was obtained. We emphasise that this comparison is only for illustrative purposes.

Our estimates of the detection parameters also match prior knowledge: a previous playback experiment for the same kiwi species reported effective detection radius of about 300-400 m, or an area of about 25 ha for a single recorder and software processing (Digby et al., 2013).

One major assumption limiting the use of SCR models with automated call recognisers – at least in their current state (Knight et al., 2017; Priyadarshani et al., 2018b) – is that there are no false positive annotations. To minimise their rate in this study, all calls suggested by the detection software were manually reviewed. This way, instead of trying to optimise the balance of false negatives and false positives, which can have unpredictable effects on the SCR models, the user needs to balance false negatives vs. the review effort – a tradeoff that can be solved with the framework presented here. While we cannot reliably quantify the number of false positives that remained after the review, we expect it to be very few, given the distinctive pattern of the kiwi call. Misidentification would be a much bigger issue for other bird species, and also amphibian sounds (McClintock et al., 2010; Mortimer and Greene, 2017).

As better recognisers become available, we expect the number of detections that need human review to decrease; this will further shift the balance in favor of automated processing methods. In the current paper, we chose to use a basic wavelet recogniser, in order to give a conservative estimate of the improvements brought by automation. More complex machine learning models are frequently used for bird call recognition, and may produce fewer false positives (Stowell et al., 2018). However, for kiwi call recognition in long-term recordings, an extension of the wavelet method used here has been shown to perform well (Priyadarshani et al., 2020). Ongoing developments are combining the detections into a smaller number of segments to review, and focusing user effort on the least certain annotations to further reduce the occurrence of false positives.

An interesting alternative was proposed by Barré et al. (2019), who used all detected cues (including false positives) in their models of bat activity, and afterwards tested the conclusions for robustness. Unfortunately, the measurement error added this way would be a particular problem for monitoring rare species, and those vocalising at lower range, where more environmental noise is present (Priyadarshani et al., 2018b). Nonetheless, a metric that allows quantifying this problem would be a valuable extension to our framework.

Although we demonstrated the issues in evaluating two particular types of survey – occupancy models and grid-based SCR – various other study designs are common in ecoacoustics, and can be evaluated using similar ideas, providing they result in a single estimate of abundance and its precision. Soundscape studies (Pijanowski et al., 2011; Sueur et al., 2014), for example, use acoustic indices to characterise a community directly, without estimating the abundance of each species. Recently, Bradfer-Lawrence et al. (2019) showed how a similar framework, focusing on the overall precision of the index estimation, can be used to optimise the design of soundscape studies. Evaluating surveys of species richness has been discussed in Darras et al. (2016) and Darras et al. (2018), although their focus was on estimating bias, not efficiency. Even if we consider only single-species, single-population surveys using autonomous recorders, different configurations of the recorders are possible, leading to different design issues.

The overlapping grid configuration used in this study is uncommon: in a recent survey, Sugai et al. (2019) found that 71% of passive acoustic monitoring studies used only a single recorder per site. However, many different designs, including single-recorder ones, can be expressed as special cases of SCR, providing that some source of information about the detection function is available, such as the estimated distance to each cue for distance sampling (Borchers et al., 2015). As an example, Sebastián-González et al. (2018) estimated the relationship between distance and recorded power of a bird call in a separate experiment, and then used this function to allow distance sampling from single-recorder sites. Yip et al. (2019) showed that such a procedure results in more accurate estimation of the true density than traditional distance sampling by a human observer. Another option is to calibrate the detection function using playback experiments (Llusia et al., 2011; Darras et al., 2016; Hagens et al., 2018), although this is not trivial because it requires knowing the true loudness at which the animal vocalises naturally. In any case, the framework presented here can be used with such surveys, and even allows a comparison of them with other designs, e.g., to evaluate the cost-benefit tradeoff of using a recorder grid. The latter task would require measuring the uncertainty in the pilot estimates of the detection function; incorporating that into the current approach could be a useful future development.

Occupancy studies avoid many of these issues and ultimately may be easier to conduct in practice. The popularity and ease of use of the occupancy framework also means that more data is available for historical comparisons, and it will likely remain a common approach in monitoring. However, guiding occupancy studies by their precision relies on knowing study-specific parameters such as the true occupancy that are typically unavailable at the onset of a study. In contrast, final precision of SCR allows the experimenter to identify universally optimal design choices, which is necessary for developing methods such as call detection software. The SCR framework can be readily extended to incorporate other types of information available to the researcher, e.g., time of arrival, telemetry data, or spatial covariates (Borchers et al., 2015; Linden et al., 2018), and applied to any vocalising species. Because of this flexibility, a large variety of management decisions can be evaluated following the approach presented here. We hope that the established SCR methods, combined with our baseline results and field dataset, which allows the testing of new analysis methods on standardised data, will provide a consistent way to evaluate new developments in acoustic survey design and analysis.

## Acknowledgements

This research is supported by the New Zealand Marsden Fund, which is administered by the Royal Society of New Zealand Te Apārangi under grants (17-MAU-154) and (17-UOA-295). We also acknowledge funding from the National Science Challenge on Science for Technological Innovation, Te Pūnaha Matatini, the New Zealand Centre of Research Excellence in Complex Systems, and MBIE Security for iconic species: Kiwi Rescue managed by Manaaki Whenua - Landcare Research.

We thank Zealandia, particularly Danielle Shanahan for allowing us to position recorders in the sanctuary, and Stephen Hartley, Paul Teal, and Heiko Wittmer for lending us acoustic recorders. We are grateful to Nirosha Priyadarshani, Virginia Listanti and Rebecca Huistra for their help in the field and in annotating the recordings.

