## Supplemental Material for "Precision as a metric for acoustic survey design using occupancy or spatial capture-recapture"

Julius Juodakis[1]*, Isabel Castro[2], Stephen Marsland[1]

[1] School of Mathematics and Statistics, Victoria University of Wellington, Wellington, New Zealand

[2] Wildlife and Ecology Group, Massey University, Palmerston North, New Zealand

### Supplementary Figures and Tables

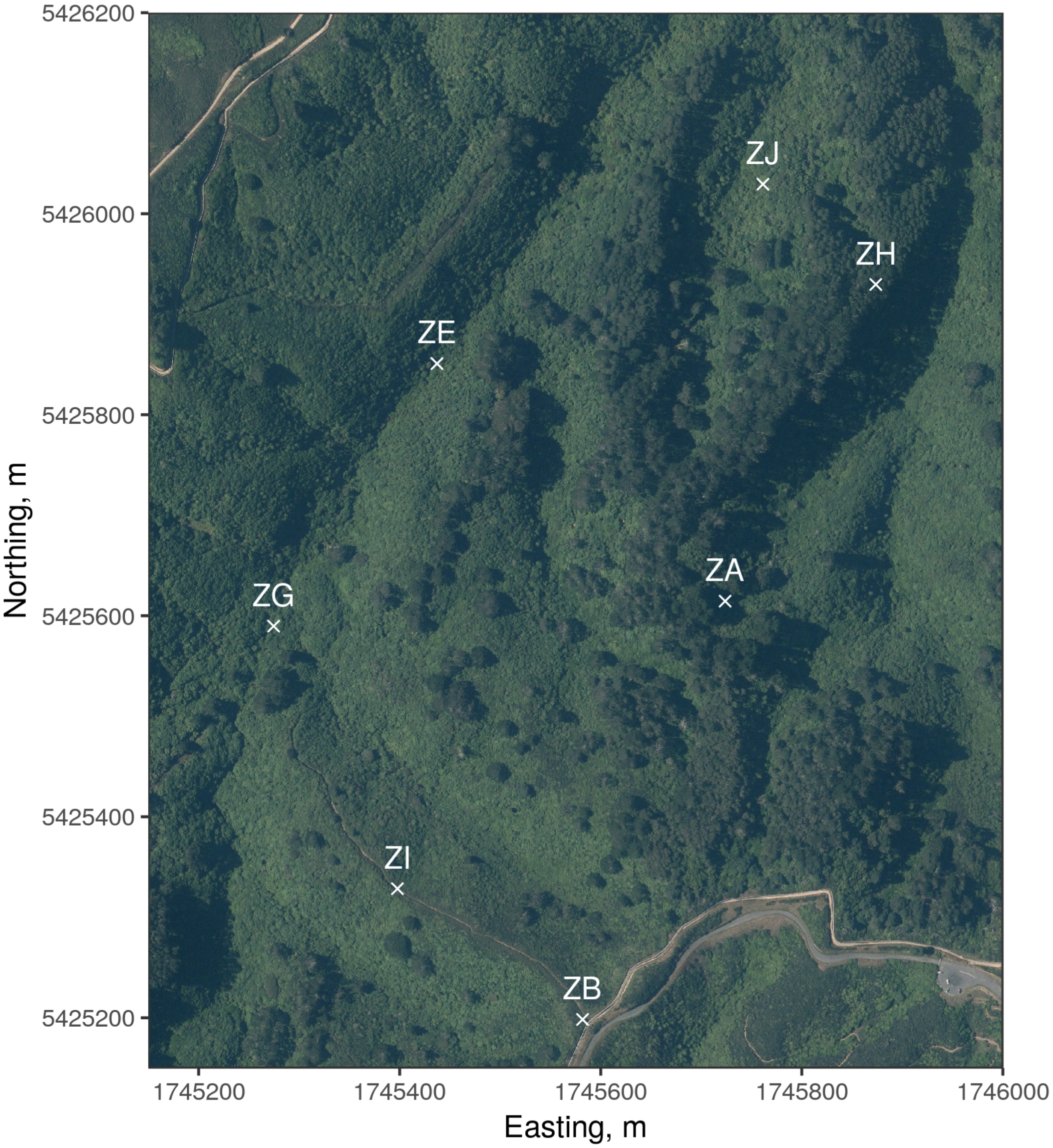


**Figure S1:** Positions of the 7 recorders used in this study within Zealandia wildlife sanctuary in Wellington, New Zealand. Northing and easting coordinates obtained by UTM projection, zone 60 South. Satellite photography sourced from the LINZ Data Service.

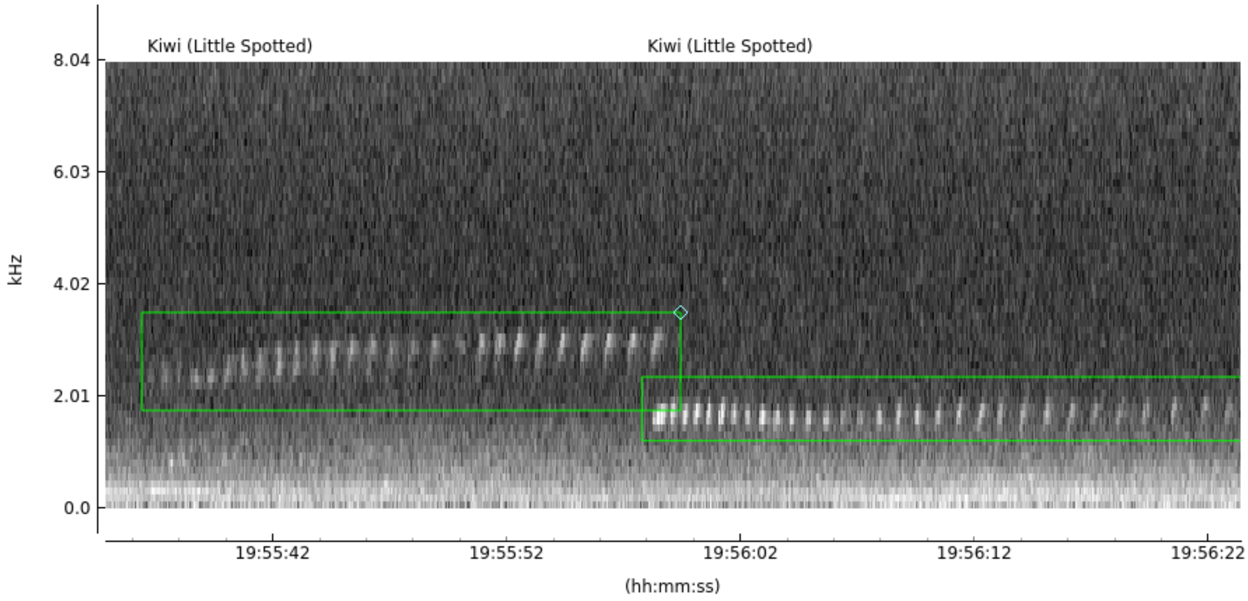


**Figure S2:** Spectrogram representation of two kiwi pukupuku calls – male followed by female. Spectrogram and annotations produced using the AviaNZ software.

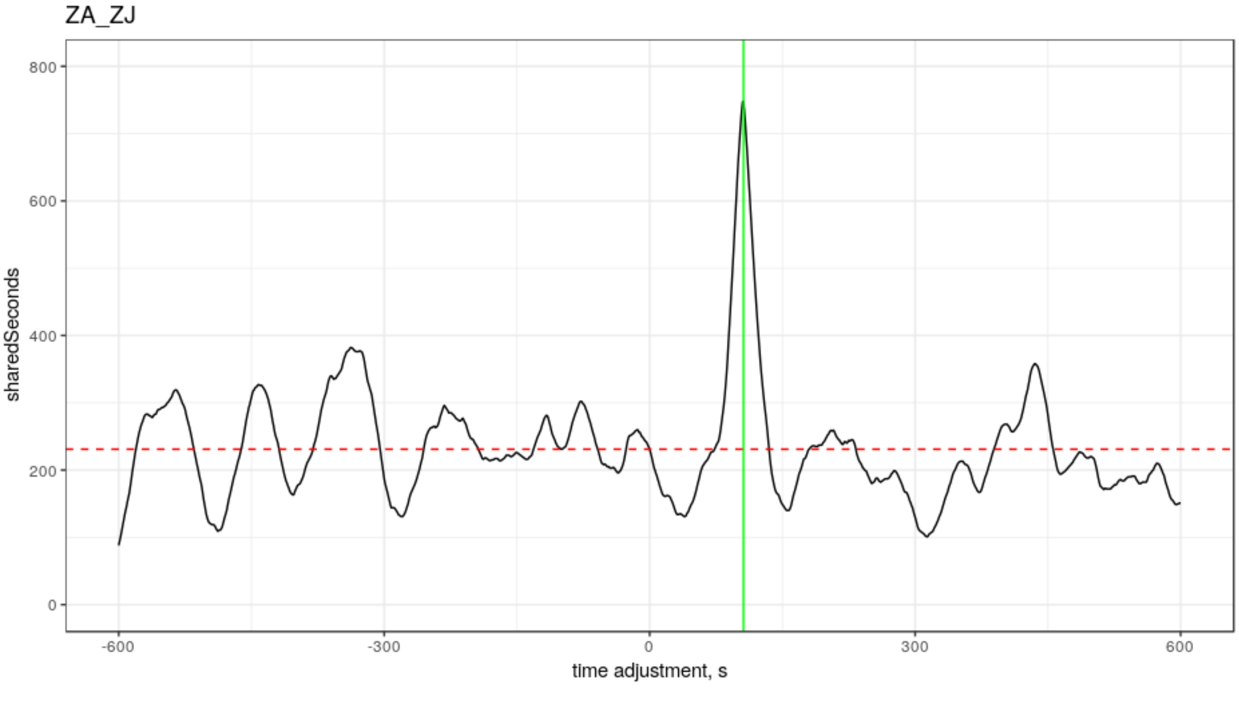


**Figure S3:** Total duration of overlapping calls across ZA-ZJ pair of recorders for various clock shifts. ZA clock is shifted between 600 s lag (left) to 600 s lead (right). Clear peak is identified, marked with a green line. Call overlap without clock adjustment (i.e. at adjustment 0) is marked with a red dashed line.

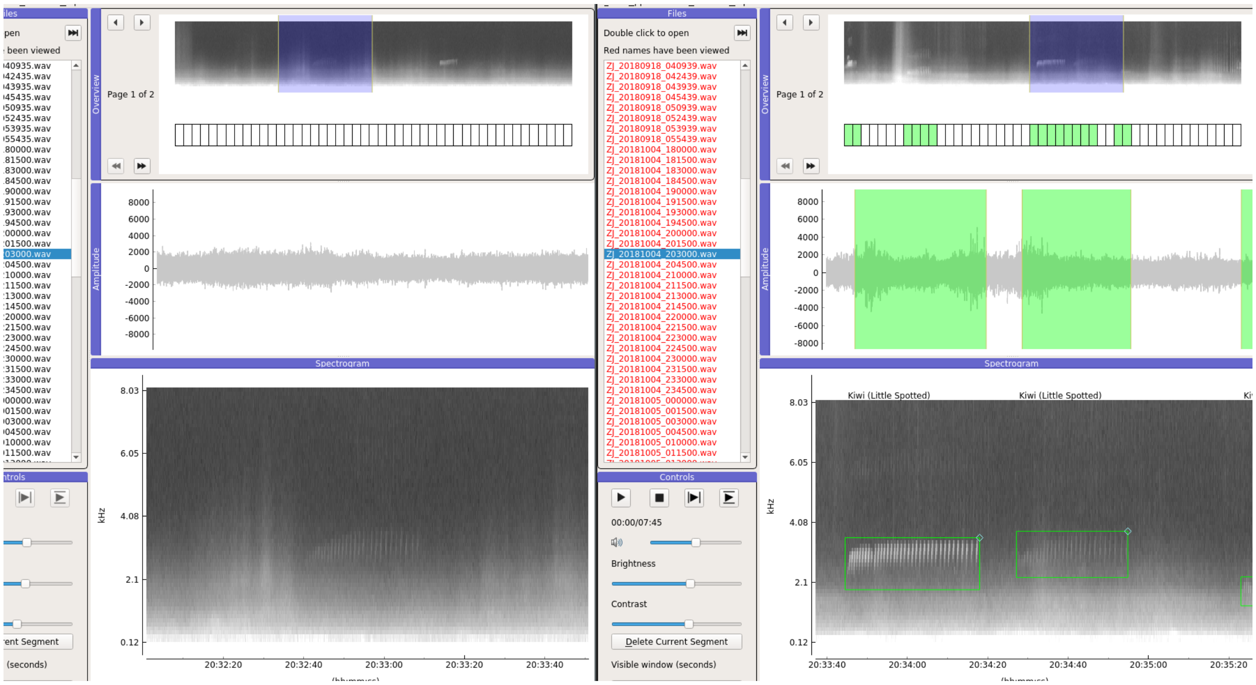


**Figure S4:** Example of overlapping calls as seen in AviaNZ. Spectrogram positions aligned based on identified clock lag (about 100 s lag on ZA). Both ZA (left) and ZJ (right) recorders capture the same call around 20:34:40 on ZJ. Call at 20:34:00 appears loud on ZJ, but not detected on ZA, indicating that it is close to the ZJ recorder.

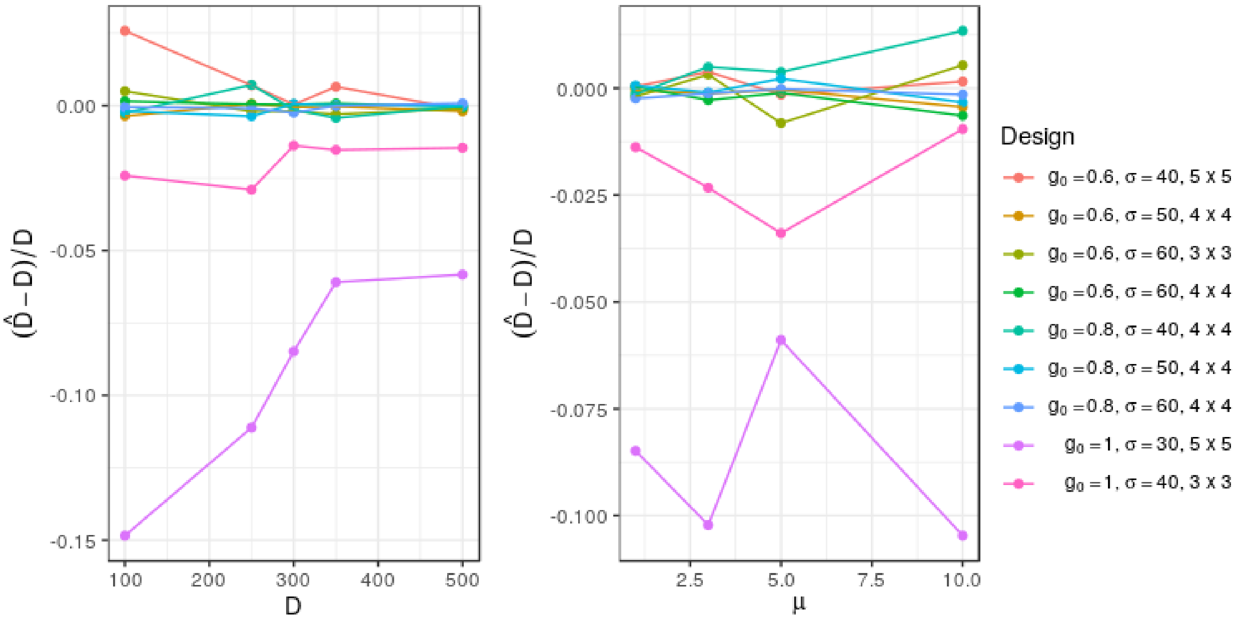


**Figure S5:** Relative bias of SCR estimates from different study designs, obtained from simulations. Call locations were simulated to be independent with density $D$ (left), or dependent, replicated at the same location $\mu$ times, with overall density of 300 (right). For each design setting (colors), 100 datasets were simulated.

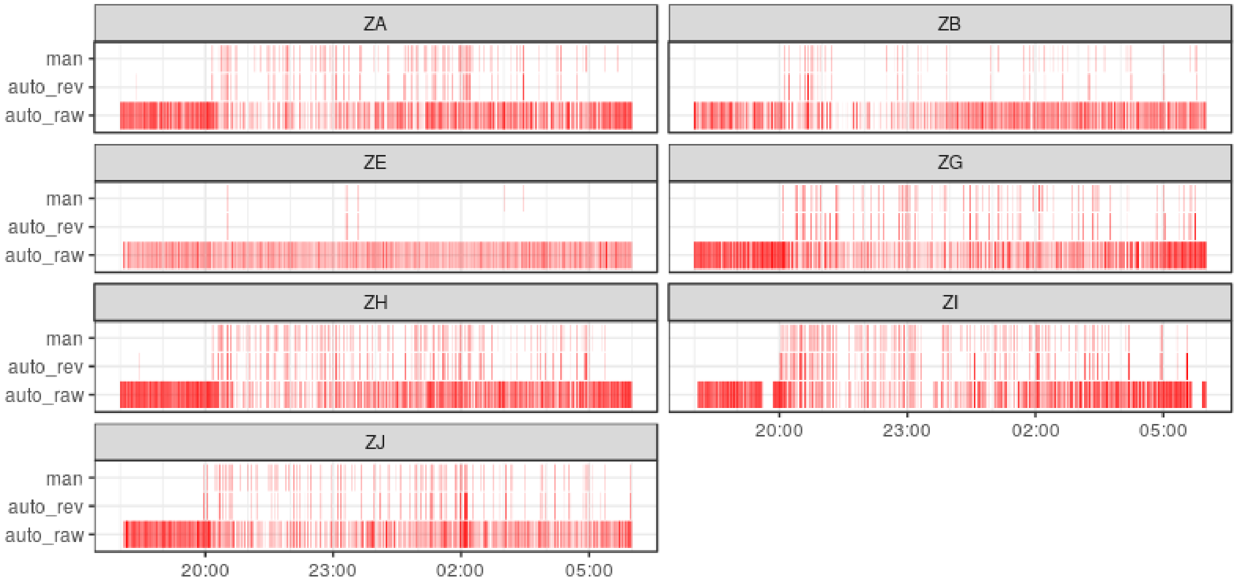


**Figure S6:** Automated annotations before review (auto_raw), automated annotations after review (auto_rev), and manual annotations (man) for Oct 6 night across 7 recorders. Red segments correspond to sections of audio marked as calls.

| Recorder | Sensitivity | Precision | Specificity | Accuracy |
|---|---|---|---|---|
| ZA | 59.67% | 90.38% | 99.60% | 97.26% |
| ZB | 62.07% | 79.25% | 99.59% | 98.68% |
| ZE | 65.93% | 91.75% | 99.98% | 99.88% |
| ZG | 72.85% | 82.97% | 99.18% | 97.80% |
| ZH | 67.42% | 86.48% | 99.10% | 96.61% |
| ZI | 73.63% | 78.95% | 98.23% | 96.20% |
| ZJ | 72.91% | 83.16% | 99.08% | 97.53% |

**Table S1:** Concordance between manual and automatic (human-reviewed) annotations for Oct 6 18:00-Oct 7 06:00 for the 7 recorders. Concordance measures (sensitivity, false positive rate, precision, specificity, and accuracy) calculated following standard definitions, after discretizing annotations into 1 second intervals.

| Recorders / annotations | Sensitivity per 15 s | Specificity per 15 s | Accuracy per 15 s |
|---|---|---|---|
| all / manual | 74.87% | 99.09% | 95.38% |
| -ZA -ZI / manual | 74.79% | 99.23% | 96.06% |
| -ZB -ZH / manual | 76.84% | 99.02% | 95.67% |

**Table S2:** Changes in concordance and SCR measures using all 7 recorders, excluding ZA and ZI, or excluding ZB and ZH, based on Oct 6 night data. Sensitivity, specificity, accuracy obtained by comparing manual and automatic annotations over 15 second blocks: presence of call (automatically or manually annotated) in a 15 s block was considered as positive for that detection method.
